# Multiplexed targeted resequencing identifies coding and regulatory variation underlying phenotypic extremes of HDL-cholesterol in humans

**DOI:** 10.1101/235887

**Authors:** Sumeet A. Khetarpal, Paul L. Babb, Wei Zhao, William F. Hancock-Cerutti, Christopher D. Brown, Daniel J. Rader, Benjamin F. Voight

## Abstract

Genome-wide association studies have uncovered common variants at many loci influencing human complex traits and diseases, such as high-density lipoprotein cholesterol (HDL-C). However, the contribution of the identified genes is difficult to ascertain from current efforts interrogating common variants with small effects. Thus, there is a pressing need for scalable, cost-effective strategies for uncovering causal variants, many of which may be rare and noncoding. Here, we used a multiplexed inversion probe (MIP) target capture approach to resequence both coding and regulatory regions at seven HDL-C associated loci in 797 individuals with extremely high HDL-C vs. 735 low-to-normal HDL-C controls. Our targets included protein-coding regions of *GALNT2, APOA5, APOC3, SCARB1, CCDC92, ZNF664, CETP*, and *LIPG* (>9 kb), and proximate noncoding regulatory features (>42 kb). Exome-wide genotyping in 1,114 of the 1,532 participants yielded a >90% genotyping concordance rate with MIP-identified variants in ~90% of participants. This approach rediscovered nearly all established GWAS associations in *GALNT2, CETP*, and *LIPG* loci with significant and concordant associations with HDL-C from our phenotypic-extremes design at 0.1% of the sample size of lipid GWAS studies. In addition, we identified a novel, rare, *CETP* noncoding variant enriched in the extreme high HDL-C group (P<0.01, Score Test). Our targeted resequencing of individuals at the HDL-C phenotypic extremes offers a novel, efficient, and cost-effective approach for identifying rare coding and noncoding variation differences in extreme phenotypes and supports the rationale for applying this methodology to uncover rare variation—particularly non-coding variation--underlying myriad complex traits.

## Introduction

While genome-wide association studies (GWAS) have elucidated the role of common genetic variation to many human complex traits and diseases, the role of rare genetic variation in complex traits remains poorly defined [1]. This is especially true for rare noncoding variants, which are not captured by whole exome sequencing (WES) currently being applied to large numbers of participants. Blood lipid levels are among the most heritable biomarkers of disease risk and protection [2]. One strategy to capture novel variation that may include putatively causal variants is targeted resequencing of genes at candidate loci for lipid traits. Indeed, this approach has been applied to the follow-up of initial GWAS studies for low-density lipoprotein cholesterol (LDL-C) and triglycerides (TG) [3–5]. These efforts have largely sequenced the coding regions of candidate genes, with the goal of identifying protein-altering variants that may have a profound functional impact. However, given that the majority of GWAS-implicated variants are in the noncoding genome [6,7] the contribution of rare noncoding variants to these traits is underexplored.

Plasma levels of high density lipoprotein cholesterol (HDL-C) are highly heritable. There are >70 loci significantly associated with HDL-C levels through testing of common variants (minor allele frequency, MAF > 0.05) on genome-wide genotyping arrays [8,9]. However, pinpointing the causal variants and genes from these associated loci is challenging. Current efforts to resolve this have included fine mapping of identified loci to determine causal variants [10,11], but these methods are limited in that they focus on common single nucleotide polymorphisms (SNPs) with generally small effect sizes. Given that common SNPs are estimated to explain only a fraction of the heritability of HDL-C levels [8], additional variance may be explained by low frequency (MAF = 0.01-0. 05) and/or rare variation (MAF < 0.01) not yet captured in existing genotyping arrays and imputation reference panels. Furthermore, the identification of rare, causal, noncoding variants with strong effect sizes on HDL-C may help to delineate causal and heritable mechanisms governing HDL metabolism that could directly relate to CHD risk. One limitation hampering targeted sequencing efforts for the noncoding genome is the relatively poor annotation of functional elements most likely to harbor variants of significance. A related issue is that targeted sequencing efforts are costly and scale with the size of the genomic targets, so methods have largely been developed for reliably amplifying and sequencing coding regions of genes. Thus, there is a pressing need for efficient and scalable method for capturing the noncoding genome to apply to large populations to uncover causal variation underlying complex traits such as HDL-C.

Here, we investigated the feasibility of targeting the noncoding regions of candidate gene loci to identify rare variants that differ in frequency at extremes of HDL-C levels using a cost-effective approach that could be extended to larger numbers of samples. We adapt a recently reported target capture method involving **<u>M</u>**olecular **<u>I</u>**nversion **<u>P</u>**robes (MIPs) [12,13] for amplifying genomic targets utilized for autism spectrum disorder candidate gene sequencing. We performed targeted resequencing of seven HDL loci including both coding and noncoding regions in a cohort of 1,532 subjects with either extremely high or low HDL-C, and show the ability to capture noncoding regions of the genome using this method. Our results validate previously reported coding and noncoding SNP associations with HDL-C, identify gene-level associations in these seven regions with this trait, and also show the promise of large-scale targeted resequencing of noncoding regions for complex traits.

## Results

### Candidate regions for targeted sequencing

We sought to develop an approach for multiplexed targeted sequencing that could identify noncoding variants, uncover novel noncoding variation in HDL-C candidate genes could underlying phenotypic extremes, and test the hypothesis that noncoding variation at these loci could contribute significantly to these extreme phenotypes in a manner similar to that of coding variants traditionally identified by targeted resequencing approaches to date. Thus, we performed a targeted resequencing study of HDL candidate gene regions in 1,532 participants with either extremely high HDL-C (mean plasma HDL-C of 107 mg/dL, >95^th^ percentile for age and sex, 797 participants) vs. low HDL-C controls (plasma HDL-C between 20 mg/dL and 25^th^ percentile for age and sex, 735 participants; **Table 1**).

**Table 1.** Characteristics of participants for MIP targeted sequencing.

|  | <u>High HDL Cohort</u> |  |  | <u>Low HDL Cohort</u> |  |  | High vs. Low HDL Cohort (T-test) |
| --- | --- | --- | --- | --- | --- | --- | --- |
|  | All (n=789) | Males (n=228) | Females (n=561) | All (n=743) | Males (n=454) | Females (n=289) |  |
| **Age** (SD) | 58 (13) | 59 (15) | 58 (12) | 55 (13) | 56 (12) | 53 (15) | P<0.0001 |
| **Caucasian** (%) | 86.2 | 89.9 | 84.7 | 61.5 | 65.0 | 56.1 | N/A |
| **Ashkenazi** (%) | 7.9 | 8.3 | 7.7 | 2.6 | 3.5 | 1.0 | N/A |
| **Black** (%) | 4.6 | 2.2 | 5.5 | 27.5 | 23.3 | 33.9 | N/A |
| **Total Cholesterol** (mg/dL) | 240 (42) | 227 (40) | 245 (42) | 177 (72) | 172 (74) | 185 (68) | P<0.0001 |
| **HDL-C** (mg/dL) | 107 (21) | 94 (19) | 112 (19) | 32 (11) | 31 (12) | 34 (8) | P<0.0001 |
| **LDL-C** (mg/dL) | 127 (60) | 127 (40) | 127 (71) | 100 (59) | 96 (58) | 105 (61) | N.S. |
| **TG** (mg/dL) | 77 (34) | 78 (37) | 77 (32) | 266 (566) | 270 (537) | 259 (610) | P<0.0001 |

Participants were recruited from the Penn High HDL Study as previously described. All lipid measurements were performed on plasma collected after participants fasted overnight. Comparisons of absolute measurements were performed using a Student's unpaired T-test of all High HDL Cohort participants vs. all Low HDL Cohort participants. All absolute data is reported as mean ± S.D.

We selected seven candidate loci for targeted sequencing of coding and noncoding regions in our cohorts (**Figure 1**). Four of the targeted loci, *APOC3, SCARB1, CETP*, and *LIPG* have known roles in HDL metabolism for which loss-of-function has been shown to elevate HDL-C in humans [14]. To explore the hypothesis that rare noncoding variants may underlie GWAS-implicated loci for HDL-C levels, we selected three HDL-C loci newly identified through GWAS, *GALNT2, SBNO1*, and the *CCDC92-ZNF664* region for our targeted sequencing. Some sequencing efforts have suggested that *GALNT2* coding variants segregate with elevated HDL-C while a recent report from our group found an opposite result for two rare coding variants [15]. Similarly, the contribution of either coding or noncoding rare variants at the *CCDC92-ZNF664* and *SBNO1* loci to HDL metabolism remains completely unexplored. Therefore, we evaluated these loci for rare coding and noncoding variants to better determine the directional relationship of these genes with HDL-C beyond the initial common variant associations.

**Figure 1.**
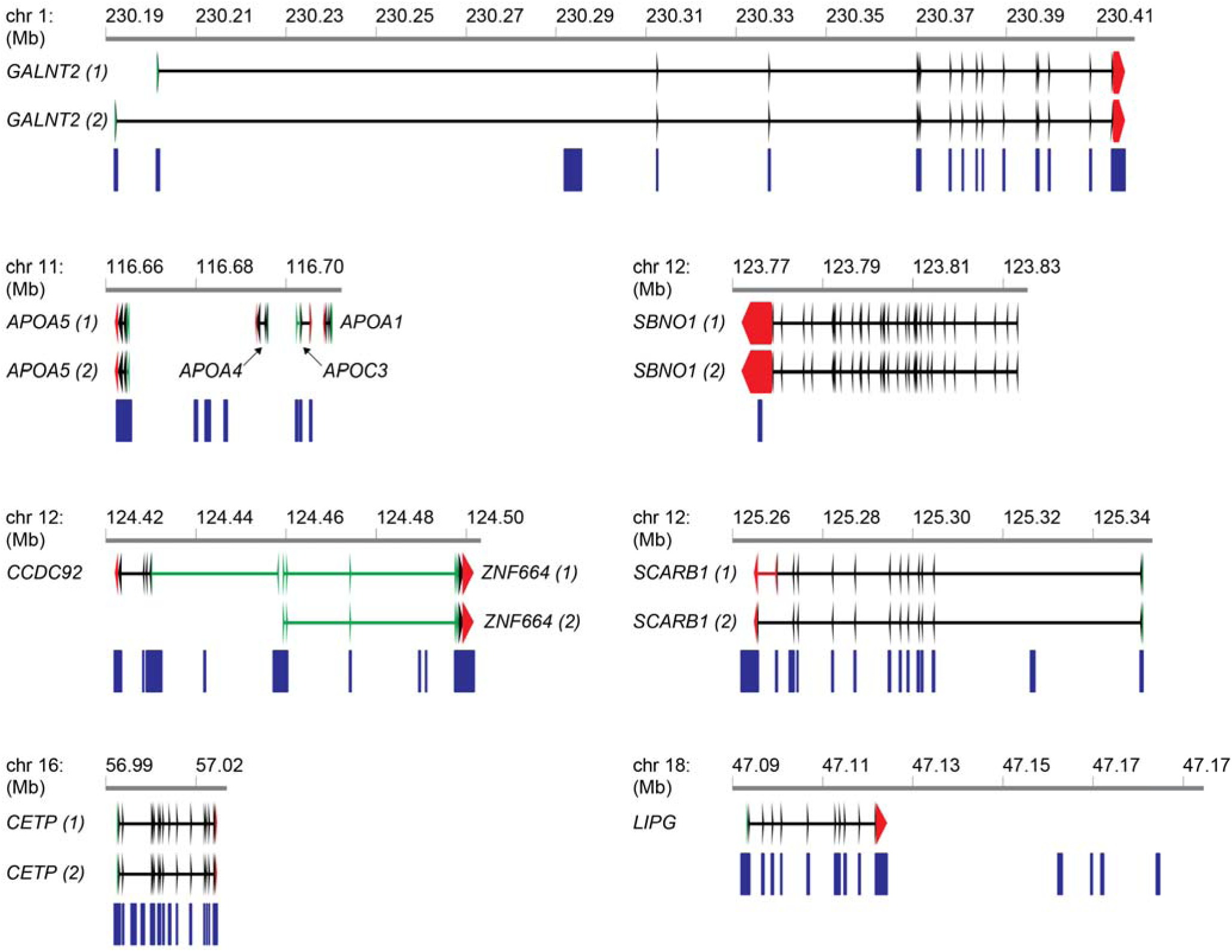
Candidate gene regions for MIP targeted sequencing. All coordinates correspond to genomic build GRC37/hg19. Blue boxes correspond to MIP target locations.

### Variants identified by MIP sequencing

We performed multiplexed capture of the genomic targets using 569 MIPs in 1,532 participants and sequenced all samples together after dual-index barcoding on the Illumina HiSeq2500 sequencing platform (**Supplementary Figure 1**). Genomic targets included 9,075 bp of protein coding sequence, 31,371 bp of noncoding UTR and intronic sequence, and 10,874 bp of noncoding intergenic sequence for a total target footprint of 51,320 bp (see **Materials and Methods**). Multiplexed sequencing across the 1,532 samples resulted in a median sequencing coverage of 110-fold per base from a single HiSeq2500 sequencing run. We observed a high uniformity of target coverage per MIP across the subjects in our cohort, with approximately 489 MIPs (86%) demonstrating coverage of >10-fold depth in each sequenced participant.

Following sequence read quality control, reads were aligned on an individual sample basis, and the alignments were then merged for joint genotyping (**Materials and Methods**). Raw variant calls were hard filtered based on alignment metrics, and then subjected to secondary variant-level and sample-level quality control pipelines to remove any additional outliers (**Supplementary Figure 2**). Next, filtered samples underwent principal component analysis to inspect for any cryptic population structure present in our cohort, identify any individual outlier samples, examine any clustering of MIP capture batches, and visualize demographic relationships in the context of 1000 Genomes samples and variants (Phase 3 version 5a; **Supplementary Figures 3-7**). After filtering MIP samples on the basis of these criteria, a total of 1500 out of 1532 original samples remained for further variant analysis.

To validate the variants identified from our MIP sequencing, we genotyped 1,114 of the 1,532 participants (681 high HDL-C individuals and 433 low HDL-C individuals) on the probe-based Illumina Exome Array [16]. Among the variants genotyped on this array, 38 were within our target regions. We observed a high concordance rate in variant discovery between MIP sequencing and genotyping results, with 32 of 38 SNPs overlapping on the Exome Array called with >90% concordance across all participants, and 987 of 1114 participants demonstrating >90% concordance of all genotyped SNPs (**Figure 2**).

**Figure 2.**
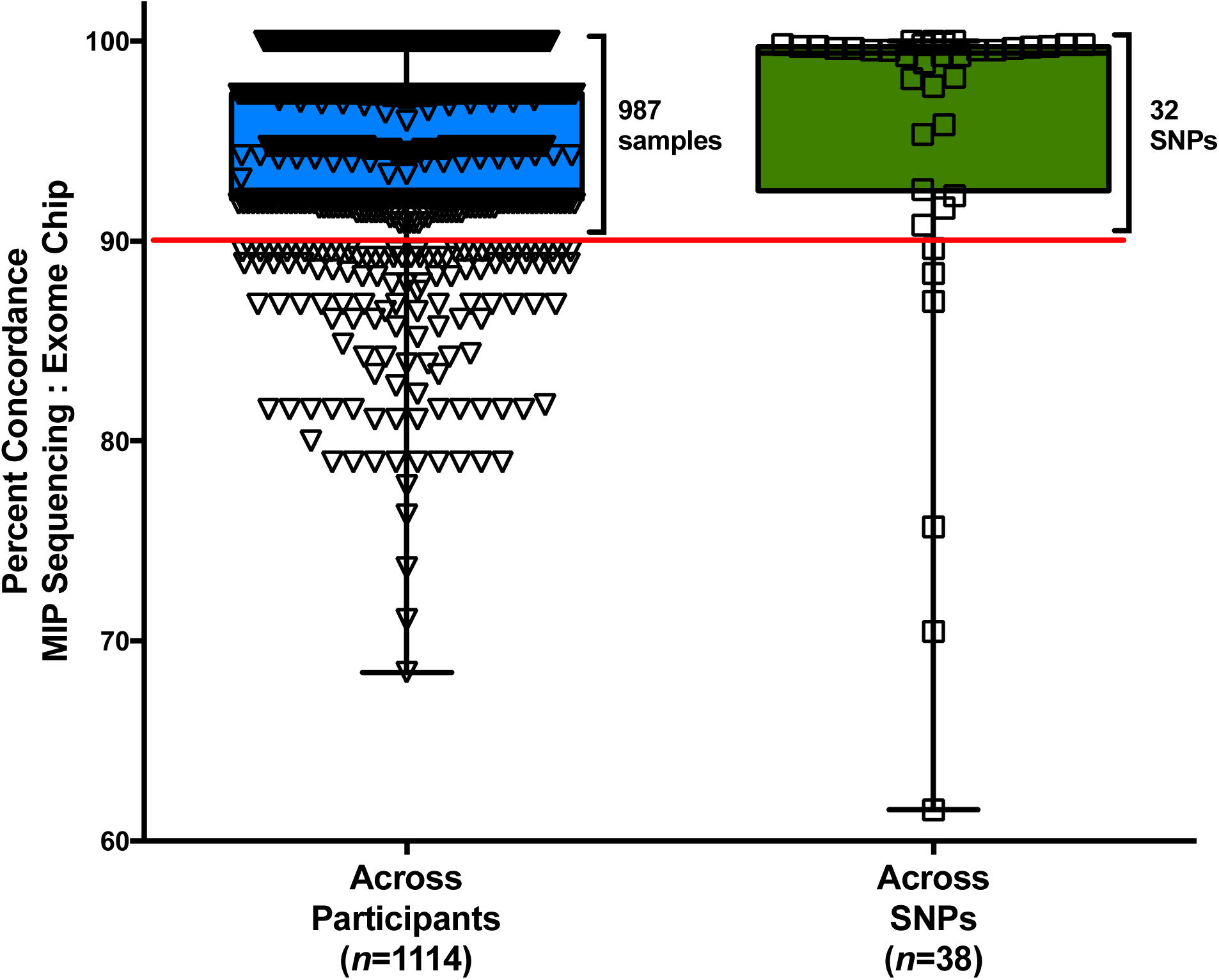
Concordance of variants identified from MIP sequencing with Exome Chip genotyping. Single nucleotide variants identified in the targeted regions by MIP-sequencing were compared to the discovery of those variants by genotyping on the Exome Chip in a subset of 1,114 participants who were included in both variant discovery efforts. A total of 38 SNPs that were included in the Exome Chip were found to overlap the targeted regions by MIPs. Box plot on the left shows the percentage of total SNPs that were found by both discovery methods for each individual (*n*=1,114 participants). Box plot on the right shows the percentage of individuals for which a given SNP was found to be concordant across the two discovery methods *n*=38 SNPs). Red line indicates those samples (left) and SNPs (right) for which concordance between MIP sequencing and the Exome Chip genotyping was >90%.

The final MIP sequencing variant call set contained 1956 SNPs and 689 distinct insertion/deletion events (indels; 78 insertions and 611 deletions) for a total of 2645 unique variants in 1500 samples. Of these, 556 correspond with previously reported variants in dbSNP (v141), suggesting that the remaining 2089 were novel discoveries without any previous annotation (**Table 2**). We also compared the frequency of identified variants across our genomic targets based on their annotated genomic position and effect on gene function (e.g. coding nonsynonymous, noncoding 5□UTR) and compared the total proportion of variants identified for a given annotation with the total amount of genomic sequence corresponding to that annotation. We found that the number of variants identified for a given annotation was proportional to the amount of sequence for a given annotation comprising the genomic target. This suggests that our MIP sequencing capture approach did not preferentially identify variants of a given annotation across our selected genomic targets (**Supplementary Figure 8**). Following quality control, genotype validation, and annotation distributions, the MIP sequencing variants were then tested for association with HDL-C using a framework sensitive to minor allele frequency (MAF) and protein coding status of the different variants (**Supplementary Figure 9**).

**Table 2.** Variants identified by MIP sequencing of 1500 extreme HDL-C participants.

| Chrom. | Genic Region | Target Size (bp) | SNPs |  | INDELs |  | Total Variants |
| --- | --- | --- | --- | --- | --- | --- | --- |
|  |  |  | Common + Low Freq. | Rare | Common + Low Freq. | Rare |  |
| 1 | *GALNT2* | 9,636 | 63 | 271 | 14 | 110 | **458** |
| 11 | *APOA5-APOC3* | 6,550 | 44 | 274 | 6 | 98 | **422** |
| 12 | *SBNO1* | 530 | 7 | 33 | - | 6 | **46** |
| 12 | *CCDC92-ZNF664* | 11,955 | 49 | 355 | 4 | 130 | **538** |
| 12 | *SCARB1* | 7,815 | 38 | 247 | 7 | 102 | **394** |
| 16 | *CETP* | 5,739 | 37 | 202 | 6 | 87 | **332** |
| 18 | *LIPG* | 9,095 | 56 | 280 | 5 | 114 | **455** |
|  | **Total** | **51,320** | **294** | **1662** | **42** | **647** | **2645** |

| Common + Low Frequency Variants (MAF≥0.01) | | | | | | |
| --- | --- | --- | --- | --- | --- | --- |
| Chrom. | Genic Region | Known<sup>a</sup> Coding | Known<sup>a</sup> Noncoding | Novel<sup>b</sup> Coding | Novel<sup>b</sup> Noncoding | Significant HDL-C associations (Score test)<sup>c</sup> |
| 1 | *GALNT2* | 4 | 56 | - | 17 | - |
| 11 | *APOA5-APOC3* | 4 | 32 | 3 | 11 | - |
| 12 | *SBNO1* | - | 2 | - | 5 | - |
| 12 | *CCDC92-ZNF664* | 4 | 36 | 5 | 8 | 9 |
| 12 | *SCARB1* | 6 | 26 | 1 | 12 | - |
| 16 | *CETP* | 5 | 32 | 2 | 4 | 20 |
| 18 | *LIPG* | 4 | 45 | - | 12 | 5 |
|  | **Total** | **27** | **229** | **11** | **69** | **34** |

| Rare Variants (MAF<0.01) |  |  |  |  |  |  |
| --- | --- | --- | --- | --- | --- | --- |
| Chrom. | Genic Region | Known<sup>a</sup> Coding | Known<sup>a</sup> Noncoding | Novel<sup>b</sup> Coding | Novel<sup>b</sup> Noncoding | Significant HDL-C associations (Score test)<sup>d</sup> |
| 1 | *GALNT2* | 10 | 44 | 37 | 290 | - |
| 11 | *APOA5-APOC3* | 8 | 26 | 52 | 286 | - |
| 12 | *SBNO1* | - | 3 | - | 36 | - |
| 12 | *CCDC92-ZNF664* | 10 | 45 | 66 | 364 | - |
| 12 | *SCARB1* | 13 | 30 | 45 | 261 | - |
| 16 | *CETP* | 18 | 26 | 42 | 203 | 2 |
| 18 | *LIPG* | 8 | 59 | 44 | 283 | - |
|  | **Total** | **67** | **233** | **286** | **1723** | **2** |

<sup>a</sup> “Known” variants were those for which an rsID existed in dbSNP (v141), or were able to be ascertained in publically available variant databases including 1000 Genomes,
the NHLBI Exome Variant Server and the Exome Aggregation Consortium (ExAC) database.
<sup>b</sup> “Novel” variants were all other variants not listed as “Known” above.
<sup>c</sup> Number of single variant associations with HDL-C using the Score test [76], at or below the experimental significance threshold of $P < 1.49 \times 10^{-4}$ (testing only 336 common and low frequency variants).
<sup>d</sup> Number of single variant associations with HDL-C using the Score test [76], at or below the experimental significance threshold of $P < 1.89 \times 10^{-5}$ (testing all 2645 variants).
Single nucleotide variants (SNPs) and insertion-deletion variants (INDELs) were assessed for each gene region (GRC37/hg19) and were processed using sample-level and variant-level quality control filters (**Materials and Methods**). Minor alleles of identified variants were compared for frequency in the high vs. low HDL cohort by the Score test statistic. Noncoding variants included any variants that were not present in protein-coding regions of the gene regions, including splice-site, intronic, 5' UTR, 3' UTR and intergenic variants.

### Association of single variants from targeted sequencing with extremely high HDL-C

We tested the association of 336 common and low frequency (MAF ≥ 0.01) SNPs and indels identified with high vs. low HDL levels, and observed 34 alleles at significantly greater frequencies among the high HDL-C participants (P < 1.49 x 10^−4^, Score test, **Table 3**). Of these, 17 were previously reported by the Global Lipids Genetics Consortium GWAS study [8].

**Table 3.** Significant single variant associations with high HDL-C.

| Region | Chrom. | Position | Variant | dbSNP<br>rsID | Type | Variant<br>Call Rate | MAF<sup>a</sup> | † | Score<br>Statistic | Score<br>P-value<sup>b</sup> |
| --- | --- | --- | --- | --- | --- | --- | --- | --- | --- | --- |
| *CCDC92-*<br>*ZNF664* | 12 | 124421453 | T/C | rs9863 | noncoding | 0.9993 | 0.4109 | † | 4.0051 | 6.20E-05 |
|  | 12 | 124427306 | T/A | rs11057401 | coding | 1 | 0.3407 | † | 4.7169 | 2.40E-06 |
|  | 12 | 124428162 | T/A | rs4930725 | noncoding | 0.9987 | 0.3632 | † | 4.2263 | 2.38E-05 |
|  | 12 | 124428331 | T/C | rs4930726 | noncoding | 0.9873 | 0.3754 | † | 4.4037 | 1.06E-05 |
|  | 12 | 124429279 | G/A | rs3186071 | noncoding | 0.9973 | 0.3259 | † | 4.1498 | 3.33E-05 |
|  | 12 | 124430612 | G/A | rs4765305 | noncoding | 0.9660 | 0.4824 | † | 4.1397 | 3.48E-05 |
|  | 12 | 124430812 | G/A | rs4765335 | noncoding | 0.9953 | 0.3985 | † | 4.0566 | 4.98E-05 |
|  | 12 | 124431049 | G/A | rs11835839 | noncoding | 0.9740 | 0.4182 | † | 4.8946 | 9.85E-07 |
|  | 12 | 124499839 | C/T | rs3768 | noncoding | 0.9993 | 0.2255 | † | 3.9722 | 7.12E-05 |
| *CETP* | 16 | 56995236 | C/A | rs1800775 | noncoding | 0.8893 | 0.3212 | † | 7.3626 | 1.80E-13 |
|  | 16 | 56995814 | G/A | rs34498052 | noncoding | 0.9580 | 0.0010 |  | 5.3163 | 1.06E-07 |
|  | 16 | 56996158 | T/C | rs3816117 | noncoding | 0.9920 | 0.4755 | † | 9.7746 | 1.45E-22 |
|  | 16 | 56996211 | G/A | rs711752 | noncoding | 0.9880 | 0.4295 | † | 7.8694 | 3.56E-15 |
|  | 16 | 56996288 | G/A | rs708272 | noncoding | 0.9887 | 0.4413 | † | 7.6112 | 2.72E-14 |
|  | 16 | 56998918 | A/G | rs12720926 | noncoding | 0.9360 | 0.3650 | † | 7.8158 | 5.46E-15 |
|  | 16 | 56999258 | A/C | rs7203984 | noncoding | 0.9747 | 0.2309 | † | -8.3195 | 8.83E-17 |
|  | 16 | 56999328 | C/T | rs11508026 | noncoding | 0.9873 | 0.3964 | † | 9.4414 | 3.68E-21 |
|  | 16 | 57001254 | T/TCACA | rs12720908 | noncoding | 0.9780 | 0.1953 | † | -7.8050 | 5.95E-15 |
|  | 16 | 57001274 | AC/A | rs200751500 | noncoding | 0.8853 | 0.1325 | † | 5.9512 | 2.66E-09 |
|  | 16 | 57001438 | G/A | rs12444012 | noncoding | 0.2433 | 0.4932 | † | 4.5664 | 4.96E-06 |
|  | 16 | 57004889 | G/A | rs7205804 | noncoding | 0.9753 | 0.3568 | † | 6.9836 | 2.88E-12 |
|  | 16 | 57005301 | C/T | rs1532625 | noncoding | 0.9840 | 0.3581 | † | 8.2715 | 1.32E-16 |
|  | 16 | 57005883 | G/A | rs374409989 | noncoding | 0.8733 | 0.0023 |  | 5.3838 | 7.29E-08 |
|  | 16 | 57007353 | C/T | rs5883 | coding | 0.9847 | 0.0735 | † | 5.4895 | 4.03E-08 |
|  | 16 | 57007446 | T/G | rs11076176 | noncoding | 0.9940 | 0.1851 | † | -6.8759 | 6.16E-12 |
|  | 16 | 57015091 | G/C | rs5880 (Ala390Pro) | coding | 1 | 0.0350 | † | -4.9197 | 8.67E-07 |
|  | 16 | 57016092 | G/A | rs5882 | coding | 0.9973 | 0.3737 | † | -4.9708 | 6.67E-07 |
|  | 16 | 57017319 | G/A | rs1800777 (Arg468Gln) | coding | 0.9973 | 0.0247 | † | -5.4589 | 4.79E-08 |
|  | 16 | 57017474 | G/A | rs289741 | noncoding | 0.9347 | 0.3574 | † | -5.2733 | 1.34E-07 |
|  | 16 | 57017662 | G/A | rs1801706 | noncoding | 0.9913 | 0.1725 | † | 4.8147 | 1.47E-06 |
|  | 16 | 57017796 | G/A | rs289743 | noncoding | 0.9440 | 0.2256 | † | -3.8050 | 1.42E-04 |
| LIPG | 18 | 47096016 | G/A | rs1320700 | noncoding | 0.9693 | 0.2775 | † | 4.1477 | 3.36E-05 |
|  | 18 | 47158186 | T/C | rs10438978 | noncoding | 1 | 0.1920 | † | 4.7214 | 2.34E-06 |
|  | 18 | 47158234 | C/T | rs9304381 | noncoding | 1 | 0.1767 | † | 4.6760 | 2.92E-06 |
|  | 18 | 47167214 | T/C | rs4939883 | noncoding | 1 | 0.2073 | † | 4.6702 | 3.01E-06 |
|  | 18 | 47179516 | G/A | rs1943973 | noncoding | 0.9947 | 0.1079 | † | 3.8348 | 1.26E-04 |

<sup>a</sup> Common and low frequency variants with a minor allele frequency (MAF) > 0.01 are marked with a cross (†), while rare variants are not marked.
<sup>b</sup> Single variant associations with HDL-C using the Score test [76]. Only variants with P-values below experimental significance threshold of $P < 1.49 \times 10^{-4}$ are shown.
Variants (SNPs and INDELs) across targets were compared for frequency of the minor allele in high vs. low HDL participants by Score test statistic. Score test P-values where $P < 0.01$ were considered nominally statistically significant, whereas P-values below $1.27 \times 10^{-5}$ were considered to exceed the experimental significance threshold accounting for all 2645 variants called in this study. MAF refers to minor allele frequency within the sequencing cohort. Call rate refers to the fraction of 1,500 samples for which a particular variant position was sequenced and passed sample-level and variant-level quality filtering.

### Replication of HDL-C associations from GWAS through MIP sequencing

In addition to rare, noncoding variants identified from MIP sequencing, we also recovered common variants previously associated with HDL-C through the Global Lipids Genetics Consortium + MetaboChip (GLGC) GWAS [8]. In the GLGC study, 49 variants that exceeded genome-wide significance (P < 5 x 10^−8^) in their associations with HDL-C are located in regions that overlap with MIP sequencing targets. We observed all of the 49 variants in the MIP sequencing variant call set, and likewise observed all of them at common or low frequencies (MAF > 0.01) in the 1500 samples. A total of 17 of the 49 exceeded an experimental statistical threshold (Score test P < 1.49 x 10^−4^), with an additional 10 that were nominally significant (Score test P < 0.01, **Table 4, Supplementary Figures 10-14**). All of the experiment-wide significant and nominally significant associations we identified were directionally consistent with prior reports of SNPs as those loci with HDL-C levels and with comparable minor allele frequencies (MAF) to those reported for each variant from 1000 Genomes Project (Phase 3 v5a, European sample set) [17,18].

**Table 4:** Replication of GWAS-significant HDL-C associations with MIP sequencing.

| Region | Chrom. | Position | Variant | dbSNP<br>rsID | GLGC +<br>MetaboChip | 1000<br>Genomes | MIP Sequencing |  |  |  |  | * |
| --- | --- | --- | --- | --- | --- | --- | --- | --- | --- | --- | --- | --- |
|  |  |  |  |  | P-value | MAF (EUR) | Variant<br>Call Rate | MAF<br>High HDL<br>Cohort | MAF<br>Low HDL<br>Cohort | Score<br>Statistic | Score<br>P-value<sup>a</sup> |  |
| GALNT2 | 1 | 230294715 | C/A | rs4846913 | 1.98E-26 | 0.5844 | 0.9880 | 0.5962 | 0.4645 | 1.7010 | 0.0889 |  |
|  | 1 | 230294916 | C/T | rs2144300 | 4.00E-40 | 0.5844 | 0.9780 | 0.6367 | 0.5085 | 0.0547 | 0.9564 |  |
|  | 1 | 230295245 | C/T | rs12065546 | 1.50E-15 | 0.8443 | 0.9807 | 0.8773 | 0.8463 | 2.7556 | 0.0059 | * |
|  | 1 | 230295307 | C/G | rs17315646 | 1.35E-36 | 0.5844 | 0.9993 | 0.5774 | 0.4482 | 2.2484 | 0.0246 |  |
|  | 1 | 230295691 | G/A | rs4846914 | 3.51E-41 | 0.5844 | 0.9927 | 0.5893 | 0.4564 | 2.0532 | 0.0401 |  |
|  | 1 | 230295789 | A/T | rs10127775 | 7.64E-35 | 0.5844 | 0.9120 | 0.5681 | 0.4406 | 1.5240 | 0.1275 |  |
|  | 1 | 230296153 | C/T | rs10864726 | 8.65E-24 | 0.5858 | 0.9940 | 0.5878 | 0.4460 | 2.6625 | 0.0078 | * |
|  | 1 | 230296469 | AC/A | rs200933185 | 5.69E-10 | - | 0.7953 | 0.1834 | 0.1661 | 1.0820 | 0.2793 |  |
| APOA5-APOC3 | 11 | 116660686 | G/A | rs2266788 | 1.19E-35 | 0.9090 | 0.8913 | 0.9297 | 0.9201 | 1.2854 | 0.1987 |  |
|  | 11 | 116660813 | G/A | rs619054 | 3.65E-23 | 0.2230 | 0.9593 | 0.2553 | 0.1887 | 2.4642 | 0.0137 |  |
|  | 11 | 116661826 | T/C | rs2072560 | 1.13E-23 | 0.9195 | 0.8640 | 0.9137 | 0.8987 | 1.8017 | 0.0716 |  |
|  | 11 | 116662331 | G/T | rs12287066 | 1.08E-20 | 0.9420 | 0.9987 | 0.0613 | 0.1051 | -2.9418 | 0.0033 | * |
|  | 11 | 116662407 | G/C | rs3135506 | 7.74E-16 | 0.9433 | 0.8920 | 0.0561 | 0.1040 | -3.5221 | 0.0004 | * |
|  | 11 | 116662579 | C/T | rs651821 | 7.72E-26 | 0.9195 | 0.9887 | 0.9320 | 0.8720 | 3.3135 | 0.0009 | * |
|  | 11 | 116663596 | C/T | rs34003087 | 1.07E-08 | 0.0541 | 0.9967 | 0.0614 | 0.0443 | 1.4969 | 0.1344 |  |
|  | 11 | 116663707 | G/A | rs662799 | 4.16E-37 | 0.9195 | 0.9987 | 0.9257 | 0.8805 | 3.0942 | 0.0020 | * |
| CCDC92<br>-ZNF664 | 12 | 124427306 | T/A | rs11057401 | 4.53E-08 | - | 1 | 0.3821 | 0.2963 | 4.7169 | 2.40E-06 | ** |
|  | 12 | 124428331 | T/C | rs4930726 | 1.53E-09 | 0.3641 | 0.9873 | 0.3977 | 0.3515 | 4.4037 | 1.06E-05 | ** |
| SCARB1 | 12 | 125259888 | A/G | rs838876 | 7.33E-33 | 0.3259 | 0.9980 | 0.6141 | 0.6172 | -2.1569 | 0.0310 |  |
|  | 12 | 125260645 | A/G | rs838878 | 3.96E-30 | 0.3100 | 1 | 0.6508 | 0.5995 | 0.2166 | 0.8286 |  |
|  | 12 | 125261441 | G/A | rs838879 | 9.37E-33 | 0.3100 | 0.3713 | 0.5610 | 0.4630 | 1.2783 | 0.2011 |  |
|  | 12 | 125261593 | C/T | rs838880 | 6.38E-32 | 0.3259 | 0.9967 | 0.6066 | 0.5700 | -1.3465 | 0.1781 |  |
|  | 12 | 125261797 | G/A | rs838881 | 1.81E-31 | 0.3087 | 0.9813 | 0.6173 | 0.5785 | -0.1513 | 0.8797 |  |
|  | 12 | 125261813 | C/T | rs838882 | 1.63E-32 | 0.3087 | 0.9807 | 0.6075 | 0.5603 | 0.2010 | 0.8407 |  |
|  | 12 | 125261839 | T/C | rs838883 | 5.10E-11 | 0.0950 | 0.9867 | 0.9451 | 0.9371 | 0.7610 | 0.4466 |  |
| CETP | 16 | 56995236 | C/A | rs1800775 | 3.33E-644 | 0.4802 | 0.8893 | 0.7587 | 0.6003 | 7.3626 | 1.80E-13 | ** |
|  | 16 | 56996211 | G/A | rs711752 | 1.287E-641 | 0.4222 | 0.9880 | 0.5267 | 0.3252 | 7.8694 | 3.56E-15 | ** |
|  | 16 | 56999258 | A/C | rs7203984 | 3.59E-517 | 0.7770 | 0.9747 | 0.1199 | 0.3494 | -8.3195 | 8.83E-17 | ** |
|  | 16 | 56999328 | C/T | rs11508026 | 2.63E-318 | 0.4142 | 0.9873 | 0.5177 | 0.2677 | 9.4414 | 3.68E-21 | ** |
|  | 16 | 57004889 | G/A | rs7205804 | 5.27E-675 | 0.4235 | 0.9753 | 0.4625 | 0.2465 | 6.9836 | 2.88E-12 | ** |
|  | 16 | 57005301 | C/T | rs1532625 | 2.25E-397 | 0.4235 | 0.9840 | 0.4599 | 0.2497 | 8.2715 | 1.32E-16 | ** |
|  | 16 | 57007353 | C/T | rs5883 | 1.76E-31 | 0.0607 | 0.9847 | 0.0961 | 0.0492 | 5.4895 | 4.03E-08 | ** |
|  | 16 | 57015091 | G/C | rs5880 | 1.37E-233 | 0.9406 | 1 | 0.0193 | 0.0518 | -4.9197 | 8.67E-07 | ** |
|  | 16 | 57016092 | G/A | rs5882 | 2.21E-58 | 0.3325 | 0.9973 | 0.6063 | 0.6479 | -4.9708 | 6.67E-07 | ** |
|  | 16 | 57017474 | G/A | rs289741 | 7.64E-160 | 0.3219 | 0.9347 | 0.6115 | 0.6757 | -5.2733 | 1.34E-07 | ** |
|  | 16 | 57017662 | G/A | rs1801706 | 1.09E-15 | 0.1939 | 0.9913 | 0.2114 | 0.1314 | 4.8147 | 1.47E-06 | ** |
|  | 16 | 57017762 | C/G | rs289742 | 1.97E-61 | - | 0.9413 | 0.8819 | 0.8741 | -2.0851 | 0.0371 |  |
| LIPG | 18 | 47093790 | C/T | rs2000812 | 2.02E-08 | 0.7744 | 0.9833 | 0.8211 | 0.8420 | 0.4481 | 0.6541 |  |
|  | 18 | 47093864 | C/T | rs2000813 | 1.08E-23 | 0.2902 | 0.9993 | 0.3155 | 0.2231 | 2.9368 | 0.0033 | * |
|  | 18 | 47118219 | T/C | rs3786248 | 2.85E-14 | 0.0660 | 0.9780 | 0.0577 | 0.0305 | 1.7461 | 0.0808 |  |
|  | 18 | 47118398 | T/C | rs9958734 | 1.52E-13 | 0.0660 | 0.9627 | 0.0737 | 0.0626 | 1.2914 | 0.1966 |  |
|  | 18 | 47157400 | T/C | rs2000825 | 2.93E-24 | 0.8245 | 0.9967 | 0.8763 | 0.8354 | 3.3554 | 0.0008 | * |
|  | 18 | 47158186 | T/C | rs10438978 | 1.56E-27 | 0.8179 | 1 | 0.8666 | 0.7452 | 4.7214 | 2.34E-06 | ** |
|  | 18 | 47158234 | C/T | rs9304381 | 3.06E-24 | 0.8193 | 1 | 0.8711 | 0.7721 | 4.6760 | 2.92E-06 | ** |
|  | 18 | 47164717 | A/G | rs7239867 | 6.22E-64 | 0.8259 | 0.9493 | 0.8507 | 0.8314 | 2.1619 | 0.0306 |  |
|  | 18 | 47164926 | T/C | rs6507937 | 6.52E-24 | 0.8259 | 0.9893 | 0.8792 | 0.8433 | 3.6424 | 0.0003 | * |
|  | 18 | 47167214 | T/C | rs4939883 | 1.80E-66 | 0.8193 | 1 | 0.8537 | 0.7272 | 4.6702 | 3.01E-06 | ** |
|  | 18 | 47167407 | T/C | rs4939884 | 5.13E-24 | 0.8206 | 0.9900 | 0.8783 | 0.8515 | 3.4407 | 0.0006 | * |
|  | 18 | 47179516 | G/A | rs1943973 | 2.75E-54 | 0.8430 | 0.9947 | 0.9105 | 0.8724 | 3.8348 | 0.0001 | ** |

<sup>a</sup> Single variant associations with HDL-C using the Score test [76]. P-values below experimental significance threshold of $P < 1.49 \times 10^{-4}$ are marked with a double asterisk (\*\*), while nominally significant P-values are marked with a single asterisk (\*).
Variants identified by MIP sequencing were compared for associations with HDL-C with the Global Lipids Genetics Consortium+MetaboChip (GLGC) GWAS results [8]. In the GLGC study, 49 variants exceeded genome-wide significant ( $P < 5 \times 10^{-8}$ ) in their associations with HDL-C are situated in regions that overlap with MIP sequencing targets. Of these 49 GLGC variants, 17 recovered by MIP sequencing exceeded the experimental threshold ( $P < 1.49 \times 10^{-4}$ ) and another 10 were nominally significant ( $P < 0.01$ ) using the Score test. In all of these cases, variants recovered by MIP sequencing displayed consistent directionality of association with HDL-C as the GLGC GWAS. The minor allele frequencies (MAF) as obtained for each GLGC variant from 1000 Genomes (European sample set) [17,18], and compared with the MAF of that variant observed in the MIP sequencing discovery cohort.

### Rare, novel, noncoding variants with nominally significant associations with HDL-C

Due to the small sample size of our study, we expected modest power to demonstrate association beyond a reasonable doubt. Thus, we examined variants that exhibited nominally significant associations (P<0.01, Score test) with elevated HDL-C, and identified 68 such SNPs and indels (**Supplementary Table 5 and Table 3**). These included 54 noncoding variants (i.e., located outside of protein-coding sequence), 11 rare (MAF≤0.01) and six low frequency variants (0.01<MAF<0.05), and eight variants not previously described in dbSNP. Of the noncoding variants identified, 12 were found to have CADD scores of 10 or more, suggestive of deleteriousness to gene expression or function (**Table S5**) [19]. We evaluated the putative impact of the noncoding variants we identified across our regions by exploring overlap between these SNPs and transcription factor binding sites and microRNA seed sites, which identified multiple common noncoding variants across our loci that overlapped such regulatory features (**Table S5**). Among the noncoding SNPs with potential functional impact on gene expression is a proximal variant 21 bp upstream of the transcription start site of *CETP*, rs34498052 (chr16:56,995,814 G>A), that was previously identified in a resequencing study of 68 genes in French Canadian myocardial infarction cases and controls. Although this variant overlaps multiple epigenetic marks from ENCODE, including CpG methylation marks in HepG2 hepatocytes and HMVEC endothelial cells, it was extremely rare (MAF=0.001, allele count [AC] = 3), which made statistical interpretation challenging, as the score test is not intended or calibrated for that end of the frequency spectrum given our sample size. More conservatively, for variants identified with greater than five allelic copies among the 1500 participants, we identified a single rare, novel, noncoding SNP in a splice region of the *CETP* gene (chr16:57,005,300 G>A) that was nominally associated with high HDL-C (P=0.009, Score Test, AC=8).

We also investigated the association of these SNPs with expression of genes as expression quantitative trait loci (eQTLs) from the Genotype-Expression (GTEx) project (**Table S5 and S6**) [20]. Analysis of eQTLs across human tissues identified 21 of the 54 noncoding SNPs with at least one significant eQTL in a human tissue. Among these are a set of noncoding SNPs at the *CCDC92* locus associated with reduced *CCDC92* expression and that of other genes in subcutaneous adipose tissues, consistent with the recent identification of a sentinel SNP at this locus in LD with our identified SNPs that was associated with CAD and also with decreased *CCDC92* expression in the same tissue [21]. As another example, we show that another set of SNPs downstream of the *LIPG* gene are associated with *LIPG* gene expression in skeletal muscle and skin tissues. These SNPs are in LD with other GWAS-implicated SNPs downstream of *LIPG* that we previously showed to reduce endothelial lipase (EL) protein levels [22]. Thus, our MIP sequencing experiment identified multiple regulatory variants underlying high HDL-C that also correlated with cis-regulatory effects on gene expression across human tissues.

### Rare variant burden associations with extremely high HDL-C

Lastly, we tested the hypothesis that the genomic regions we targeted harbor rare variants that collectively contribute to the relationship of these genes with HDL-C levels. We performed aggregate rare variant burden using a framework that categorized rare variants (MAF<0.01) on the basis of their coding status, deleteriousness, and genic region (**Table 5 and Supplementary Figure 9**). We first identified rare coding variants believed to be non-benign in their putative functional consequence (*n*=213), organized them based on their predicted impact on protein function (*e.g.: i*) disruptive, *ii*) disruptive plus missense, or *iii*) loss-of-function; see **Materials and Methods** for definitions), and then tested aggregate rare coding variant burden across all targeted genic regions for each predicted impact category. We found that for each predicted impact category the collection of all rare coding variants did not exhibit a level of rare variant burden that was significantly associated with HDL-C. Similarly, variant aggregation over the coding regions of the individual gene targets separately (*n*=8) did not identify any individual region with significant variant burden associated with high vs. low HDL-C (Collapsing test; **Table 5**).

**Table 5:** Rare variant burden test associations of MIP sequencing variants with high HDL-C.

| CODING |  |  |  |  |  |  |  |  |  |  |  |  |  |
| --- | --- | --- | --- | --- | --- | --- | --- | --- | --- | --- | --- | --- | --- |
| Chr | Genic Region | Variants | <u>Disruptive<sup>a</sup></u> |  | P-value | Variants | <u>Disruptive and missense<sup>b</sup></u> |  | P-value | Variants | <u>Loss-of-function<sup>c</sup></u> |  | P-value |
|  |  |  | □ (SE) |  |  |  | □ (SE) |  |  |  | □ (SE) |  |  |
| 1 | *GALNT2* | 15 | 0.03 (0.33) |  | 0.92 | 34 | 0.10 (0.22) |  | 0.64 | 15 | 0.03 (0.33) |  | 0.92 |
| 11 | *APOA5* | 15 | -0.36 (0.30) |  | 0.23 | 33 | -0.37 (0.23) |  | 0.10 | 14 | -0.35 (0.31) |  | 0.25 |
| 11 | *APOC3* | 5 | -0.38 (0.71) |  | 0.59 | 7 | -0.60 (0.65) |  | 0.36 | 5 | -0.38 (0.71) |  | 0.59 |
| 12 | *CCDC92* | 20 | 0.25 (0.27) |  | 0.36 | 33 | 0.00 (0.22) |  | 0.98 | 20 | 0.25 (0.27) |  | 0.36 |
| 12 | *ZNF664* | 2 | 2.11 (1.38) |  | 0.13 | 14 | 0.37 (0.32) |  | 0.25 | 2 | 2.11 (1.38) |  | 0.13 |
| 12 | *SCARB1* | 24 | 0.28 (0.27) |  | 0.29 | 37 | 0.27 (0.21) |  | 0.20 | 23 | 0.30 (0.27) |  | 0.27 |
| 16 | *CETP* | 23 | -0.10 (0.27) |  | 0.71 | 33 | -0.06 (0.22) |  | 0.77 | 12 | -0.52 (0.45) |  | 0.25 |
| 18 | *LIPG* | 14 | -0.10 (0.29) |  | 0.73 | 32 | 0.01 (0.21) |  | 0.95 | 13 | -0.12 (0.29) |  | 0.67 |
| - | All Coding<sup>d</sup> | 118 | 0.04 (0.14) |  | 0.78 | 223 | -0.09 (0.12) |  | 0.44 | 104 | 0.01 (0.14) |  | 0.95 |
| NONCODING |  |  |  |  |  |  |  |  |  |  |  |  |  |
| Chr | Genic Region | Variants | <u>Non-rare alleles at multiallelic positions retained<sup>e</sup></u> |  | P-value | Variants | <u>Non-rare alleles at multiallelic positions removed<sup>f</sup></u> |  | P-value |  |  |  |  |
|  |  |  | □ (SE) |  |  |  | □ (SE) |  |  |  |  |  |  |
| 1 | *GALNT2* | 335 | 0.04 (0.12) |  | 0.72 | 333 | 0.10 (0.12) |  | 0.40 |  |  |  |  |
| 11 | *APOA5* | 100 | -0.10 (0.14) |  | 0.51 | 100 | -0.10 (0.14) |  | 0.51 |  |  |  |  |
| 11 | *APOA5-APOC3* intergenic | 151 | 0.31 (0.12) |  | 0.009 | 149 | -0.02 (0.14) |  | 0.89 |  |  |  |  |
| 11 | *APOC3* | 62 | 0.30 (0.20) |  | 0.13 | 62 | 0.30 (0.20) |  | 0.13 |  |  |  |  |
| 12 | *SBNO1* | 39 | 0.21 (0.21) |  | 0.31 | 39 | 0.21 (0.21) |  | 0.31 |  |  |  |  |
| 12 | *CCDC92* | 251 | -0.08 (0.12) |  | 0.48 | 251 | -0.08 (0.12) |  | 0.48 |  |  |  |  |
| 12 | *ZNF664* | 156 | 0.03 (0.13) |  | 0.84 | 156 | 0.03 (0.13) |  | 0.84 |  |  |  |  |
| 12 | *SCARB1* | 292 | 0.20 (0.14) |  | 0.15 | 290 | 0.16 (0.12) |  | 0.17 |  |  |  |  |
| 16 | *CETP* | 229 | -0.06 (0.12) |  | 0.63 | 229 | -0.06 (0.12) |  | 0.63 |  |  |  |  |
| 18 | *LIPG* | 343 | -0.34 (0.14) |  | 0.02 | 341 | -0.16 (0.12) |  | 0.19 |  |  |  |  |
| - | All Noncoding | 1958 | -0.63 (0.94) |  | 0.50 | 1950 | -0.27 (0.38) |  | 0.48 |  |  |  |  |

- <sup>a</sup> Disruptive coding variants included nonsense (stop-gained), frameshift, splice-donor, splice-acceptor, stop lost, start lost, inframe insertion, and inframe deletion variants as annotated from Ensembl's Variant Effect Predictor (VEP) tool. - <sup>b</sup> Collection of disruptive (<sup>a</sup>) plus missense coding variants. Missense variants were defined as nonsynonymous amino acid-altering variants using the dbNSFP database (v2.9.1). Variants were included in this grouping if they were identified as 'deleterious' or 'damaging' by one of the five in silico prediction tools: SIFT (deleterious), PolyPhen2 HDIV ('possibly damaging' or 'probably damaging'), PolyPhen2 HVAR ('possibly damaging' or 'probably damaging'), MutationTaster, and LRT (disruptive). - <sup>c</sup> Loss-of-function variants were defined based on LoF prediction flags (HC, LC) generated by the VEP plugin LOFTEE. This set was then filtered to remove variants that were situated in unlikely open-reading-frames, single-exon genes, or had weak phylogenetic conservation scores. - <sup>d</sup> No coding regions of the *SBNO1* region were sequenced in this study. - <sup>e</sup> Aggregation of rare noncoding variants included non-rare alleles at multiallelic positions also harboring rare variants. - <sup>f</sup> Aggregation of rare noncoding variants with non-rare alleles at multiallelic positions removed.

We then asked if the burden of rare noncoding variants across all targets contributed to extremely high HDL-C. Due to the fact that a methodological framework for predicting the potential regulatory impact of noncoding variants genome-wide has yet to be widely accepted, the rare noncoding variants were not subdivided into putative functional categories like the coding variants described above. Thus, we first analyzed all rare noncoding variants as a single group, which resulted in a variant burden that was not significantly associated with high HDL-C in our cohort (P=0.5028; **Table 5**). We next grouped rare noncoding variants by physical genic region (*n*=10) and performed variant burden analyses separately on each region. This approach identified a collection of 151 rare variants in the *APOA4-APOA5* intergenic region that were nominally significantly associated with extremely high HDL-C (P=9.43 x 10^−3^, Collapsing test; **Table 5**). Within this region, we noted a collection of three different indels as multiple alternative alleles at the position chr11:116,678,249 (hg19). Of these, a rare deletion CAA>C (MAF=0.003, AC=7) exhibited nominally significant association with high HDL-C (P=0.0427, Score test). The second allele was a common deletion (MAF=0.06, AC=138) that was not associated with high HDL-C (P=0.75, Score test). The third allele was the same common (MAF=0.26, AC=605) yet previously unreported insertion of CAA>CAAA at chr11:116,678,249 that was significantly associated with high HDL-C (P=8.9x10^−4^, Score Test) in the single variant analysis.

We hypothesized that these particular common alternative alleles were driving the nominally significant rare variant burden association signal for the *APOA4-APOA5* intergenic region. To test this, we removed it (and all other non-rare variants at multiallelic sites) and reassessed rare variant burden and found a complete attenuation of the association (P=0.43; **Table 5**), thus suggesting that the originally significant association of the cluster of *APOA4-APOA5* intergenic variants with HDL-C was driven by common alleles alone.

## Discussion

Translating GWAS trait- and disease-associated common variants to *bona fide* causal variants, genes, and biological mechanisms has been a major challenge for human genetics. This is due in part to small effect sizes of GWAS variants, and thus resequencing of candidate genes at GWAS loci at the phenotypic extremes of complex traits has become a leading approach to identify rare variants with larger effects. To date, this approach has been applied to coding regions of GWAS candidate genes, yet coding variants account for only a small fraction (approximately 11%) of all variants tagged complex trait GWAS studies [23,24], underscoring the need to search the noncoding genome for rare, putatively causal variants. Here, we utilized an inexpensive, modular, and scalable targeted sequencing approach for identifying rare noncoding variants in candidate genes influencing HDL-C, a complex trait with 72 associated loci from GWAS [8]. Our proof-of-principle resequencing study of seven candidate gene regions in 797 extremely high HDL-C vs. 735 low HDL-C participants rediscovered and validated nearly all prior GWAS-implicated tag SNPs, and revealed nearly 2,000 variants in noncoding regions of targets, including rare, novel noncoding variants that were nominally associated with HDL-C in our study. As such, our findings provide one of the first applications of a multiplexed targeted resequencing study of noncoding variants across multiple loci at the phenotypic extremes of a complex trait.

We rediscovered previously implicated variants in our cohort, along with the initial discovery of a few novel candidates requiring statistical support. Most notably, we found significant or nominally significant associations for a majority (55%) of GWAS-implicated HDL-C variants overlapping our targeted regions with consistent directionality to prior associations of these variants. However, we replicated these associations at less than 1/100^th^ the cohort size of the most recent GWAS for HDL-C (188,577 participants [8], vs. 1532 participants in our study) through our phenotypic extremes-design. We also identified three rare (MAF < 0.01) or low frequency (MAF < 0.05) nonsynonymous coding variants associated with HDL-C levels with directionalities consistent with previous reports (*CETP* Ala390Pro [25], *CETP* Arg468Gln [26], and *LIPG* Asn396Ser [26–28]). Collectively, these findings support the utility of candidate gene and noncoding locus resequencing at the extremes of a continuous trait distribution to enrich for trait-associated alleles, which may allow ascertainment of genetic associations in smaller populations than historical sizes for complex trait GWAS, such as understudied ethnicities and population isolates.

Our study also has important methodological implications for future targeted resequencing efforts. To date, MIP-sequencing has been applied to targeted sequencing of coding regions of candidate genes with a sample preparation cost of less than $1 per participant [12,29,30]. Our use of MIPs to interrogate noncoding regions of HDL-C candidate genes represents one of the first applications of this methodology for regulatory DNA regions. Our sequencing efforts were completed at a comparable cost to the prior applications, with similar target-coverage depths across coding and noncoding targets. Additionally, our modified dual-barcoding approach allowed us to multiplex all 1,532 samples for sequencing in a single lane of an Illumina HiSeq2500 sequencing run with a median base coverage per participant of 110-fold; a robust depth for novel and rare variant identification at a sequencing cost of ~$2,000. Thus, our study highlights the utility of a MIP-based approach for sequencing of noncoding regions at a low per-sample cost.

Several variants identified from our study lie in regions of candidate genes for which loss-of-function variants have been shown to raise HDL-C levels in humans. Specifically, multiple noncoding variants were found in high HDL-C participants were observed in *CETP* and *LIPG*. CETP is a circulating regulator of HDL metabolism with pharmacological and genetic inactivation, including coding and noncoding variants, associated with increased HDL-C in humans [15]. Similarly, we also identified multiple rare noncoding variants in LIPG among high HDL-C subjects. *LIPG* encodes endothelial lipase, an enzyme critical to HDL catabolism for which loss-of-function genetic variants are causal contributors to elevated HDL-C in humans [15]. Here, we expanded the allelic spectrum of rare noncoding variation in these two HDL-C modulating genes contributing to high HDL-C levels in humans. In both cases, the frequency of these mutations and our limited sample size requires further analysis in follow-up cohorts to demonstrate conclusive association of these rare alleles with HDL-C.

While epidemiological findings have consistently supported an inverse association of high-density lipoprotein cholesterol (HDL-C) with CHD [31–33], the direct role that HDL-C plays in modulating CHD risk has been highly controversial. Increasing evidence over the last decade has argued against the hypothesis that simply raising serum HDL-C levels will protect against CHD [34], most directly supported by the lack of efficacy of pharmacological elevation of serum HDL-C to lower CHD risk [35–38]. Subsequently, human genetic efforts identifying low-frequency or rare coding variants in candidate loci robustly associated with HDL-C elevation have not demonstrated a reduction in the incidence of CHD or myocardial infarction [15]. Taken collectively, these studies raise basic questions regarding the causal role of HDL in CHD biology, HDL metabolism and the medical interpretation of the phenotypic extremes of the HDL-C spectrum. Elucidation of these facets of HDL biology is therefore likely to be central in determining how HDL ultimately underlies cardiovascular disease risk.

Our current study has limitations, which serve as opportunities for further study. First, our total cohort size of 1,532 participants limits both the ascertainment of the full spectrum of very rare variants that may underlie extremely high HDL-C levels as well as the power of our statistical tests of common variant association and rare variant burden. Second, our population of high and low HDL-C participants was largely of European ancestry, thus limiting our ability to extrapolate the variants discovered to other populations. Third, we employed conventional strategies for rare-variant grouping, which focused on gene-level aggregation. However, for noncoding sequences, it was not obvious which variant grouping strategy is optimally powered, which remains an open question in the field. Finally, because we selected a finite sequence of noncoding genome with genomic annotations that we believed *a priori* would be functional and lipid related (*e.g.*, enhancer marks in liver), it remains possible that rare-variant burden either exists in other sequences we did not target here.

In conclusion, our MIP-based targeted sequencing approach has demonstrated the successful capture of noncoding regions for the discovery of rare, noncoding variants associated with HDL-C in a cohort of extremely high vs. low HDL-C participants. Though efforts to better identify the spectrum of noncoding variants underlying complex traits have initiated, including denser genotyping of noncoding variants [39] and whole-genome sequencing [26], these approaches remain expensive and not readily applicable to the study of large populations or large case-control designs. Our results offer a scalable and cost-effective targeted approach that complement future, larger candidate loci resequencing efforts for the discovery of putatively causal noncoding variants. These efforts, coupled with appropriate functional investigation of identified variants for impact on gene regulation, may substantially refine the causal genes at loci implicated from GWAS studies and also help further explain the missing heritability underlying complex traits such as HDL-C.

## Materials and Methods

### Ethics statement

All human participants of this study and all analyses performed were completed following the Declaration of Helsinki [40] and were approved by the Institutional Review Board of the Perelman School of Medicine at the University of Pennsylvania and all participants provided informed consent.

### Subject selection and ascertainment

1532 participants mostly of European ancestry, with either extremely high HDL-C (>95^th^ percentile for age and sex), or low HDL-C (20 mg/dL or higher to 25^th^ percentile for age and sex) were recruited for targeted sequencing (**Table 1**). Participants were recruited as part of the University of Pennsylvania High HDL Study (HHDL), a cross-sectional study of genetic factors contributing to elevated HDL-C levels. Individuals with elevated HDL-C (>90^th^ percentile for age and gender) were identified by physician referrals or through the Hospital of the University of Pennsylvania clinical laboratory. Plasma lipids for all subjects were measured after fasting by a clinical autoanalyzer (Hitachi). HDL-C percentiles for inclusion were calculated for individuals of European ancestry from the Framingham Heart Study Offspring cohort adjusted for age and sex.

### Molecular inversion probe design

Molecular inversion probes (MIPs) were designed according to the method and pipeline previously described by O’Roak et al [12]. Briefly, MIPs capturing chosen targets were all designed using a common 30 bp linker sequence flanked by an extension arm of 16-20 bp and a ligation sequence of 20-24 bp, with a total MIP length of 70 bp. The unique arms of the MIPs that anneal to the target sequence by complementary base pairing were designed to amplify a specific 112-150 bp target region by gap-filling and circularization. After MIP capture, a PCR amplification reaction using Nextera-like (Illumina) sequencing adaptor-containing primers (Illumina) allowed amplification with the primers annealing to the 30 bp common linker sequence (**Supplementary Figure 1**). Given prior demonstration of variability in MIP capture efficiency due to properties of annealing arm base pairing with sequences adjacent to individual targets, an initial set of 549 MIPs was designed to cover all of the proposed target in 88 unique non-overlapping segments, and a pilot-phase MIP sequencing study was performed to evaluate per sample and per MIP coverage depth in an initial set of 95 DNA samples. Based on the coverage from this run, MIPs demonstrating less than 10-fold coverage per base for more than 50% of sequenced samples were redesigned and substituted in all additional runs. From this second and final pilot-phase sequencing run, 569 MIPs were included to capture the targeted regions from genomic DNA samples from the 1,532 participants.

MIPs were designed to capture the coding sequences (exons) of the following genes (GRC37/hg19 coordinates): *GALNT2* (chr1:230338882-230415202), *APOA5* (chr11:116660886-116663095), *APOC3* (chr11:116700650-116703573), *CCDC92* (chr12:124421729-124428847), *ZNF664* (chr12:124488089-124497396), *SCARB1* (chr12:125267297-125348261), *CETP* (chr16:56995891-57017572), and *LIPG* (chr18:47088681-47110124) (**Figure 1 and Supplementary Table 1**). The total protein-coding sequence captured by the 569 MIPs corresponding to these regions was 9,075 bp. Noncoding genic regions at these loci such as 5□ untranslated regions (UTRs), 3□ UTRs, and intronic sequences were likewise targeted, for a total of 31,371 bases. Noncoding sequences, including 5□UTRs, 3□UTRs, intronic sequences and other intergenic noncoding sequences proximate to these loci, were chosen if they were previously shown to harbor variants significantly associated with HDL-C (P<5 x 10^−8^; GLGC + Metabochip GWAS), and also were found to overlap DNase I hypersensitivity sites in HepG2 cells (human hepatocellular carcinoma) from the ENCODE project [41] or enhancers in HepG2 cells from the Epigenome-Roadmap project [42]. Regions with 250 bp flanking the positions harboring these elements were selected for MIP design. The total noncoding intergenic target across the loci for which MIPs were designed to capture was 10,874 bp. The entire sum of genomic territory for targeted resequencing was 51,320 bp. MIP oligonucleotides were purchased from Eurofins Genomics with high-purity salt-free purification at a scale of 50 nmol per oligonucleotide. Lyophilized MIPs were hydrated with 1x TE buffer to a concentration of 100 μM and stored at -20 °C.

### MIP capture and amplification of targeted sequences

MIP oligonucleotides were used to capture targets from genomic DNA derived from whole blood from the participants in a manner described previously. Hydrated MIP oligonucleotides were pooled together and phosphorylated with T4 polynucleotide kinase (NEB) at 37 °C for 45 min, followed by heat inactivation at 65 °C for 20 min. Phosphorylated MIPs were used to capture genomic targets by combining with genomic DNA from each participant (100 ng of each individual sample; ratio of 800:1 of each MIP copy to haploid genome copy) using NEB Hemo Klentaq (NEB) and Ampligase for 24 hrs at 60 °C in a thermocycler. 96 samples were individually captured in one reaction by individually plating reactions in a 96-well thermocycler plate. A total of 16 plates of 96 samples apiece were processed. Reactions were digested with Exonuclease I and Exonuclease III (NEB) after incubations for 45 min at 37 °C and then 2 min at 95 °C. Digested MIP capture reactions were PCR amplified using primers with barcoded adapter sequences (**Supplementary Table 2**). In order to sequence all 1532 samples from a single multiplexed pool, a dual-barcoding strategy similar to that of Illumina’s Nextera protocol was employed. To provide unique combinations of forward and reverse primers for all 1536 samples (1532 individual subjects plus four ddH_2_O controls) across the 16 plates, a common forward barcoded adapter primer was used for each plate, and 96 unique reverse barcoded adapter primers were used for each of the 96 samples within a plate. PCR reactions to ligate adapters and barcode MIP capture reactions were completed with iProof master mix reagent (Bio-Rad). Barcoded and PCR amplified MIP capture reactions were then pooled together at equal volumes, purified using AMPure magnetic bead purification (Agencourt) at 0.9-fold the total volume of the pooled reaction, and visualized on agarose gels. Purified, pooled capture reactions were then sequenced in paired-end mode (150 bp X 150 bp) on Illumina MiSeq and HiSeq2500 sequencers using standard Nextera sequencing reagents plus a custom pool of Nextera-like sequencing primers (**Supplementary Table 3**). All MIP oligonucleotides, adapters, PCR primers and sequencing primers were synthesized by Eurofins MWG Operon.

### MIP sequencing

Initial MIP sequences were obtained as paired-end FASTQ reads and generated in three separate sequencing runs (one lane of sequences from a single MiSeq run and two lanes of HiSeq2500 RapidRun from two independent runs). Coverage estimates were calculated on a per-run basis, whereas variant calling utilized reads from all three runs. De-multiplexing was performed using CASAVA v1.8.2’s bcl2fastq conversion script (Illumina), and all reads were inspected using FastQC (http://www.bioinformatics.babraham.ac.uk/projects/fastqc/) and processed using Trimmomatic v0.32 to remove adapter artifacts, sequencing artifacts, and low quality bases [43].

### Read alignment and variant calling

Sequences were aligned to the UCSC hg19 human genome build on a per-sample and per-sequencing run basis using BWA v0.7.8 (MEM algorithm) [44,45] and the resulting alignment files were compressed and sorted using SAMtools v0.1.19 [46]. The variant calling was conducted utilized Genome Analysis Toolkit v3.5 (GATK; [47]), and pre-processing of each sample’s lane-specific alignment files was performed in accordance with the established GATK’s ‘Best Practices’ workflow [48,49]. This workflow featured duplicate read removal using Picard v1.141 (Picard website: http://broadinstitute.github.io/picard), and run-specific insertion-deletion (indel) realignment and base recalibration using GATK and hg19 “Gold Standard” variant catalogs (dbSNP v138 database: http://www.ncbi.nlm.nih.gov/SNP/, [50,51]). Run-specific alignments were then merged for each sample, and subjected to a second round of indel realignment and base recalibration with GATK. Preliminary sample-level variants were called using GATK’s HaplotypeCaller tool in gVCF mode at base-pair resolution, and all known variants were annotated with their corresponding dbSNP v138 identities. Sample-level variant callsets were then combined and joint-genotyped with GATK. SNPs and indels called at this stage were evaluated using metrics collected by Picard and GATK, and then hard-filtered on the basis of variant-class-specific criteria (**Supplementary Table 4**) in order to flag potential false positives. To avoid the inclusion of soft-clipped adapter artifacts, all variants falling outside of the MIP target regions were removed using VCFtools v0.1.13 [52].

### Validation by exome array genotyping

Genomic DNA from 1,114 of the 1,500 participants whose samples passed QC were also subject to genotyping using the Exome Array (HumanExome BeadChip v1.0, Illumina, Inc., San Diego, CA). The Exome Chip contains >240,000 coding SNPs derived from all mutations found >2 times across >1 dataset among 23 separate datasets comprising a total of >12,000 individual exome and whole genome sequences. In total, 681 high HDL-C participants and 433 low HDL-C participants were genotyped using the Exome Array.

### Sample-level quality control of MIP sequencing

Quality control of samples was performed using Variant Association Tools (VAT) v2.6.1 rev2881 [53]. SNPs and indels were imported separately into VAT, sample-level and genotype-level summaries were created, and a number of filters were applied to remove outliers using VAT and VCFtools. First, any samples with a high degree of missing genotype calls (>90% variant positions) for either SNP or indel variant sets were removed. Next, any samples with mean genotype quality scores below 10 were removed. Lastly, to identify any demographic outliers or cryptic relatedness, the MIP sequencing samples were compared to samples from the 1000 Genomes Project [17,18] (*n*=2504, Phase 3 v5a). Samples from the two datasets were combined and multidimensional scaling (MDS) was performed with PLINK v1.07 [54] using only variants in regions that overlapped with MIP targets. SNP and indel genotypes were analyzed separately. After plotting principle components, any outlier MIP sequencing samples that did not cluster with the other samples were flagged, and subsequently removed from downstream analyses. After applying all of these filters, 1500 of the original 1532 MIP sequencing samples (97.9%) were retained.

### Variant-level quality control of MIP sequencing

Variant statistics, including minor allele frequency (MAF), genotype quality, call rates, novel and known variant counts, transition-transversion ratio (TS:TV), and insertion-deletion ratio were computed across the MIP sequencing cohort variant sets using VAT. Again, SNPs and indels were analyzed separately. To reduce the rate of inaccurate variant calls, any variant with a high proportion of missing genotype calls (>90%) across the filtered samples was removed, as were variants with a low maximum genotype quality scores (<10). In addition, any variants that were no longer variable following earlier sample-level filtering were also removed. After applying all of these filters, 1956 SNP variants and 689 indel variants were retained (2645 total variants).

### Variant annotation

The post-QC filtered and annotated SNP and indel call sets were then combined using VCFtools, and the union of these variants was used as input for variant annotation. Individual alleles at multiallelic sites were normalized using bcftools and then individually annotated with RefSeq gene coordinates for human genome build hg19 (RefSeq database: http://www.ncbi.nlm.nih.gov/books/NBK21091/) using bcftools v1.3.1 (http://samtools.github.io/bcftools/; [55]) to include features such as full gene lengths, protein coding sequences, exon and intron boundaries, and 5□ and 3□ UTRs. Following this, all variants were annotated using Ensembl’s Variant Effect Predictor (VEP) rel. 84 [56,57] in conjunction with the following plugins and tests to append transcript information and score the deleteriousness of different mutations: Ensembl_transcriptid, Uniprot_acc, Uniprot_id, Uniprot_aapos, SIFT_pred, Polyphen2_HDIV_pred, Polyphen2_HVAR_pred, LRT_pred, MutationTaster_pred, MutationAssessor_pred, FATHMM_pred, PROVEAN_pred, MetaSVM_pred, and MetaLR_pred [58–71]. In addition, the dbNSFP v2.9.1 [46] database plugin for VEP was used to evaluate missense (nonsynonymous) mutations, and the LOFTEE plugin [LOFTEE website: https://github.com/konradjk/loftee] was used to identify protein-truncating variants predicted to disrupt gene function on the basis of annotation details and evolutionary sequence conservation.

### Association testing and statistics

Association testing of all MIP sequencing variants was performed in the context of a framework that applied different tests on the basis of each variant’s MAF and protein-coding status (**Supplementary Figure 9**). Rare variants (MAF<0.01, *n*=1958 multiallelic alternative alleles retained, *n*=1950 with multiallelics pruned) were aggregated in different groupings that underwent rare variant burden tests, whereas common and low frequency variants (MAF≥0.01, *n*=336) were individually subjected to single variant association tests. Different experimental P-value thresholds of significance were estimated and applied depending on the particular kind of test and/or grouping of variants involved. For all association tests the HDL-C levels of the samples were treated as a dichotomous phenotype of “high” (>95^th^ percentile for age and gender) or “low” (<25^th^ percentile for age and gender). Association testing was performed using EPACTS v3.2.6 (EPACTS website: http://csg.sph.umich.edu/kang/epacts/home), and R v3.2.5 (R Core Team 2015).

Rare variant burden tests were computed after grouping variants in different aggregations based on coding status. Rare variants that were identified as protein-coding (with explicit CDS annotations, *n*=353) were grouped either together as a single group, “All coding”, or were divided up according to gene (*n*=8 groups). Each of these two aggregation strategies was then further refined to three groupings that included only coding variants that were flagged as either “Disruptive” (*n*=118), “Disruptive + Missense” (*n*=223), or “Loss-of-Function” (*n*=104). A total of 130 coding variants annotated as “benign” or “likely benign” were not tested. Following this categorization strategy, six aggregates of rare coding variants were independently tested for HDL-C association using the Collapsing burden test [72–75].

Similarly, rare variants in noncoding regions (*n*=1966 sites without specific ‘CDS’ annotations) were grouped either together as a single group denoted as “All noncoding”, or grouped by “genic region” (*n*=10 groups). These two aggregation strategies were then tested independently using the Collapsing burden test.

To account for and correct multiple testing, the total number of variant groupings within the different aggregation strategies (coding=3+24; noncoding=1+10; total grouping=38) resulted in 38 hypotheses tested. This value was then Bonferroni-corrected (□=0.05) and resulted in an experimental threshold of P=1.32 x 10^−3^ for significant associations of rare variant burden to HDL-C. Associations were considered ‘nominally significant’ with P<0.01.

Meanwhile, single variant associations for high vs. low HDL-C levels were computed for all common and low frequency variants (MAF≥0.01, *n*=336) with the Score test statistic [76]. The biological sex and self-identified ethnicity (White [non-Ashkenazi], Black, Ashkenazi) of each sample were used as phenotypic covariates in the regression analysis. Single variant associations (*n*=336 tests) were considered statistically significant if P-values for associations were below the Bonferroni-corrected (□=0.05) experimental-wide threshold of P=1.49 x 10^−4^. Single variant associations with P<0.01 were considered nominally significant. In order to investigate signals of rare variant burden for different genic regions and correct for multiallelic inflation, we also ran single variant association tests for all variants of all frequencies (*n*=2654). In this context, single variant associations (*n*=2645 tests) were considered statistically significant if P-values for associations were below the Bonferroni-corrected (□=0.05) experimental-wide threshold of P=1.89 x 10^−5^. Single variant associations with P<0.01 were considered nominally significant. In the cases of very low allele counts of extremely rare variants, these tests should be approached with caution.

## Additional Website References

Picard Tools v1.141. Available from: http://broadinstitute.github.io/picard.

Database of Single Nucleotide Polymorphisms (dbSNP). Bethesda (MD): National Center for Biotechnology Information, National Library of Medicine. (dbSNP Build ID: 138). Available from: http://www.ncbi.nlm.nih.gov/SNP/.

The NCBI handbook [Internet]. Bethesda (MD): National Library of Medicine (US), National Center for Biotechnology Information; 2002 Oct. Chapter 18, The Reference Sequence (RefSeq) Project. Available from http://www.ncbi.nlm.nih.gov/books/NBK21091/.

LOFTEE (Loss-Of-Function Transcript Effect Estimator): a plugin for the Ensembl Variant Effect Predictor (VEP) to identify LoF (loss-of-function) variation. Konrad Karczewski. Available from: https://github.com/konradjk/loftee.

EPACTS v3.2.6 (Efficient and Parallelizable Association Container Toolbox). Available from: http://csg.sph.umich.edu/kang/epacts/download/EPACTS-3.2.6.tar.gz.

R Core Team (2015). R: A language and environment for statistical computing. R Foundation for Statistical Computing, Vienna, Austria. Available from: https://www.R-project.org/.

## Supplementary Items

**Supplementary Figure File**

MIP_Supp_Figures_V8_112717.docx

**Supplementary Figure 1** Diagram of MIP target capture and sequencing reaction.

**Supplementary Figure 2** Quality control metric distributions of SNPs and INDELs called from MIP sequencing data, before and after sample-level and variant-level filtering.

**Supplementary Figure 3** Multidimensional scaling plots of quality-filtered SNPs called from MIP sequencing.

**Supplementary Figure 4** Multidimensional scaling plots of quality-filtered INDELs called from MIP sequencing.

**Supplementary Figure 5** Multidimensional scaling plots of 1000 Genomes samples using variants within MIP sequencing regions.

**Supplementary Figure 6** Multidimensional scaling plots of quality-filtered SNPs using the merged set of MIP sequencing and 1000 Genomes samples and variants.

**Supplementary Figure 7** Multidimensional scaling plots of quality-filtered INDELs using the merged set of MIP sequencing and 1000 Genomes samples and variants.

**Supplementary Figure 8** Relative proportions of variant consequence annotations of quality-filtered SNPs and INDELs called from MIP sequencing.

**Supplementary Figure 9** Association testing workflow diagram for MIP sequencing variants.

**Supplementary Figure 10** Locus plot of association signals at the *GALNT2* locus (chromosome 1) using GLGC GWAS and MIP sequencing variants.

**Supplementary Figure 11** Locus plot of association signals at the *APOA4-A5-C3-A1* locus (chromosome 11) using GLGC GWAS and MIP sequencing variants.

**Supplementary Figure 12** Locus plot of association signals at *SBNO1, CCDC92-ZNF664*, and *SCARB1* loci (chromosome 12) using GLGC GWAS and MIP sequencing variants.

**Supplementary Figure 13** Locus plot of association signals at the *CETP* locus (chromosome 16) using GLGC GWAS and MIP sequencing variants.

**Supplementary Figure 14** Locus plot of association signals at the *LIPG* locus (chromosome 18) using GLGC GWAS and MIP sequencing variants.

## Supplementary Table File

MIP_Supp_Tables_V8_121317.docx

**Supplementary Table Index**

ST_INDEX

**Supplementary Table 1** MIP sequences and coordinates.

ST_01

**Supplementary Table 2** Indexed adapter oligo sequences.

ST_02

**Supplementary Table 3** Sequencing primers.

ST_03

**Supplementary Table 4** Variant filtering criteria.

ST_04

**Supplementary Table 5** Single variant associations with high HDL-C.

ST_05

**Supplementary Table 6** Expression QTLs across human tissues from GTEx for single variants significantly associated with HDL-C.

ST_06

